# Pseudogenes confirm ongoing loss of ethylene biosynthesis in seagrasses

**DOI:** 10.64898/2026.06.09.731081

**Authors:** Sébastien Cabanac, Christophe Dunand, Catherine Mathé

## Abstract

Many flowering plant species have adopted an aquatic lifestyle, contrasting with their terrestrial ancestors. Adapting to an aquatic environment required numerous evolutionary changes, including gene expansion and contraction. One of the most striking contractions has been observed in the genomes of seagrasses, where the ACO and ACS genes, involved in ethylene biosynthesis, are very few in number or even completely absent. To confirm this adaptation, we identified traces of gene loss in the genomes of four seagrass species, in the form of pseudogenes. Surprisingly, no gene loss was found in the species that had completely lost the function of ethylene synthesis, likely indicating an ancient loss of these genes. Conversely, several pseudogenes were found in the species where the ACO and ACS genes are contracting, indicating a recent and potentially ongoing process. We used the same approach on Utricularia gibba, a submerged freshwater plant, and also found a reduced number of ACO and ACS genes. In contrast, two terrestrial species closely related to seagrasses and U. gibba found a higher number of ACO and ACS genes, with no definitive evidence of gene loss. These results confirm that the loss of ethylene biosynthesis function in seagrasses is indeed linked to gene loss and suggests that it is an adaptation to a submerged rather than a marine lifestyle.

## Introduction

Pseudogenes are sequences in the genome that share homology with functional genes, but that have lost their primary function. They can arise through mRNA retrotransposition in the case of processed pseudogenes, or through the loss of functionality of a coding gene due to a deleterious mutation. In the latter case, this represents a gene loss event, and the resulting pseudogene can be described as duplicated if it still shares homology with a functional “parent” gene, or unitary otherwise. With the increased availability of sequenced genomes, comparative genomics studies are on the rise, particularly those aimed at comparing the expansion and contraction of multigene families between species. Some examples among others include the contraction of multigene families associated with pathogenesis in *Calonectria* fungi (Rogers et al., 2022) or *Mycobacterium* bacteria (Zhu et al., 2023), or contractions associated with aquatic lifestyle in plants (Chen et al., 2026). In this type of studies, it remains difficult to determine whether a contraction actually corresponds to a loss or reduction of function in one organism, or an increase in the organism to which it is compared. Pseudogenes might help answer this question, since contractions of multigene families should leave genomic traces in the form of pseudogenes.

Among terrestrial plants, some species have re-adapted to an aquatic lifestyle, notably a large number of species belonging to the Alismatales order. These seagrass species can complete their entire life cycle fully submerged, a rare ability even among aquatic plants, which often have part of their vegetative or reproductive structures above ground (den Hartog, 1967). This adaptation to an environment very different from that of their terrestrial ancestors was accompanied by extensive gene loss and drastic morphological changes. For example, the reduction or disappearance of stomata on the leaves of aquatic plants is often observed. Even more strikingly, seagrasses appear to have a reduced number of genes involved in ethylene biosynthesis (Chen et al., 2026, 2022; Ma et al., 2024). A complete loss of the ethylene-biosynthesis-related *ACO* and *ACS* genes has even been noted in *Cymodocea nodosa* and *Zostera marina* (Ma et al., 2024). This reduction is possibly linked to the fact that this phytohormone is notably used in terrestrial plants to sense flooding (Leeggangers et al., 2023), which is not necessary in marine plants and could even lead to toxic concentration levels due to its buildup within their tissues (Ma et al., 2024).

In this article, we compared the *ACO* and *ACS* pseudogenes of four aquatic Alismatales species, *C. nodosa, Posidonia oceanica, Thalassia testudinium* and *Z. marina*, with the pseudogenes of the amphibious Alismatales *Colocasia esculenta*. Although this plant can live or survive in terrestrial, moist environments or even in shallow water, the vast majority of its shoot constantly remains at the surface. We used a similar approach on plants belonging to the dicotyledon clade, focusing on the mostly-submerged Lamiales *Utricularia gibba* and the terrestrial *Erythranthe nasuta*, because *U. gibba* is a freshwater plant and we wanted to determine whether *ACO* or *ACS* pseudogenes were found for this type of adaptation.

## Results

20315, 4449, 8406, 21147, 24630, 6941 and 16735 pseudogenes were respectively found in *C. esculenta, C. nodosa, E. nasuta, P. oceanica, T. testudinium, U. gibba* and *Z. marina*. However, many pseudogenes actually appear to be complete transposable elements, as seagrass genomes are transposon-dense (Ma et al., 2024), or unannotated functional genes. Considering only pseudogenes with strong evidences of pseudogenization (frame-shift or extra stop codon), we found 11447, 1271, 2995, 10704, 14176, 3111 and 6828 pseudogenes in *C. esculenta, C. nodosa, E. nasuta, P. oceanica, T. testudinium, U. gibba* and *Z. marina*, respectively (Table 1). Those numbers are positively correlated with genome sizes (R^2^ = 0.85).

**Table 1.** Number of duplicated, unitary, processed and uncategorized pseudogenes predicted in 5 seagrass, and 2 monocotyledons *Utricularia gibba* and *Erythranthe nasuta* genomes. Only pseudogenes with frame-shifts or more than one predicted stop codon are counted.

| Species | Total | Duplicated | Unitary | Processed | Uncategorized |
| --- | --- | --- | --- | --- | --- |
| *C. esculenta* | 11447 | 3283 | 1355 | 886 | 5923 |
| *C. nodosa* | 1271 | 271 | 291 | 119 | 590 |
| *E. nasuta* | 2995 | 675 | 640 | 183 | 1497 |
| *P. oceanica* | 10704 | 4104 | 1035 | 890 | 4675 |
| *T. testudinum* | 14176 | 5171 | 764 | 1266 | 6975 |
| *U. gibba* | 3111 | 663 | 290 | 221 | 1937 |
| *Z. marina* | 6828 | 1775 | 748 | 403 | 3902 |

We annotated the pseudogenes across the seven genomes and retrieved those predicted by homology with one of the *ACS* or *ACO* genes. We checked each pseudogene sequence to ensure it corresponded to *ACO* or *ACS* pseudogenes and verified that they were not missed exons of functional genes. We also checked the already annotated functional genes and found that four of them were truncated (*P. oceanica*’s *Posoc05g18830* and *Posoc04g07190, C. esculenta*’s *EVM0000316* and *T. testudinium*’s *Thate06g07530*). For *Posoc05g18830* and *Thate06g07530*, the missing parts of the genes were found, and *Posoc05g18830* recovered parts showed traces of pseudogenization (Table S1), but not *Thate06g07530*. We also removed four pseudogenes showing traces of intron loss or that were very similar to short fragments of a functional protein, these pseudogenes likely being retropseudogenes and therefore not true gene losses (Table S2).

In the end, we found two *ACO* pseudogenes, one in *C. nodosa* (*ψACO4*) and one in *P. oceanica* (*ψACO5*). However, the *ψACO4* from *C. nodosa* is a small fragment that could have been coded by a single exon, and it does not allow observation of intron loss or retention. It cannot therefore be confirmed whether it is actually a gene loss or a retrotransposition. We found five *ACS* pseudogenes: two *ACS7* pseudogenes in *P. oceanica* (*ψACS7_1* and *ψACS7_2*), one *ACS8* pseudogene in *P. oceanica* and in *T. testudinium* (*ψACS8*) and one *ACS2* pseudogene in *C. esculenta* (*ψACS2*). That last pseudogene is also a short fragment for which intron loss or retention cannot be confirmed (Table 2).

**Table 2.** Main characteristics of *ACO* and *ACS* pseudogenes predicted and found among genome annotations.

| Species | Pseudogene name | Completeness | Frame-shifts | In-frame stop codons | Intron-related events | “True” gene loss |
| --- | --- | --- | --- | --- | --- | --- |
| *C. nodosa* | ψACO4 | Fragment | 2 | 0 | Unknown | Unsure |
| *P. oceanica* | ψACO5 | Full | 1 | 1 | Retention | Yes |
| *C. esculenta* | ψACS2 | Fragment | 0 | 0 | Unknown | Unsure |
| *P. oceanica* | ψACS7_1 | Full | 1 | 0 | Retention | Yes |
| *P. oceanica* | ψACS7_2 | Fragment | 0 | 0 | Retention | Yes |
| *P. oceanica* | ψACS8 | Fragment | 0 | 0 | Retention | Yes |
| *T. testudinum* | ψACS8 | Fragment | 0 | 0 | Retention | Yes |

The number of *ACO* or *ACS* genes varies within species, but there is a clear trend towards contraction associated with gene loss in aquatic plants (Figure 1).

**Figure 1.**
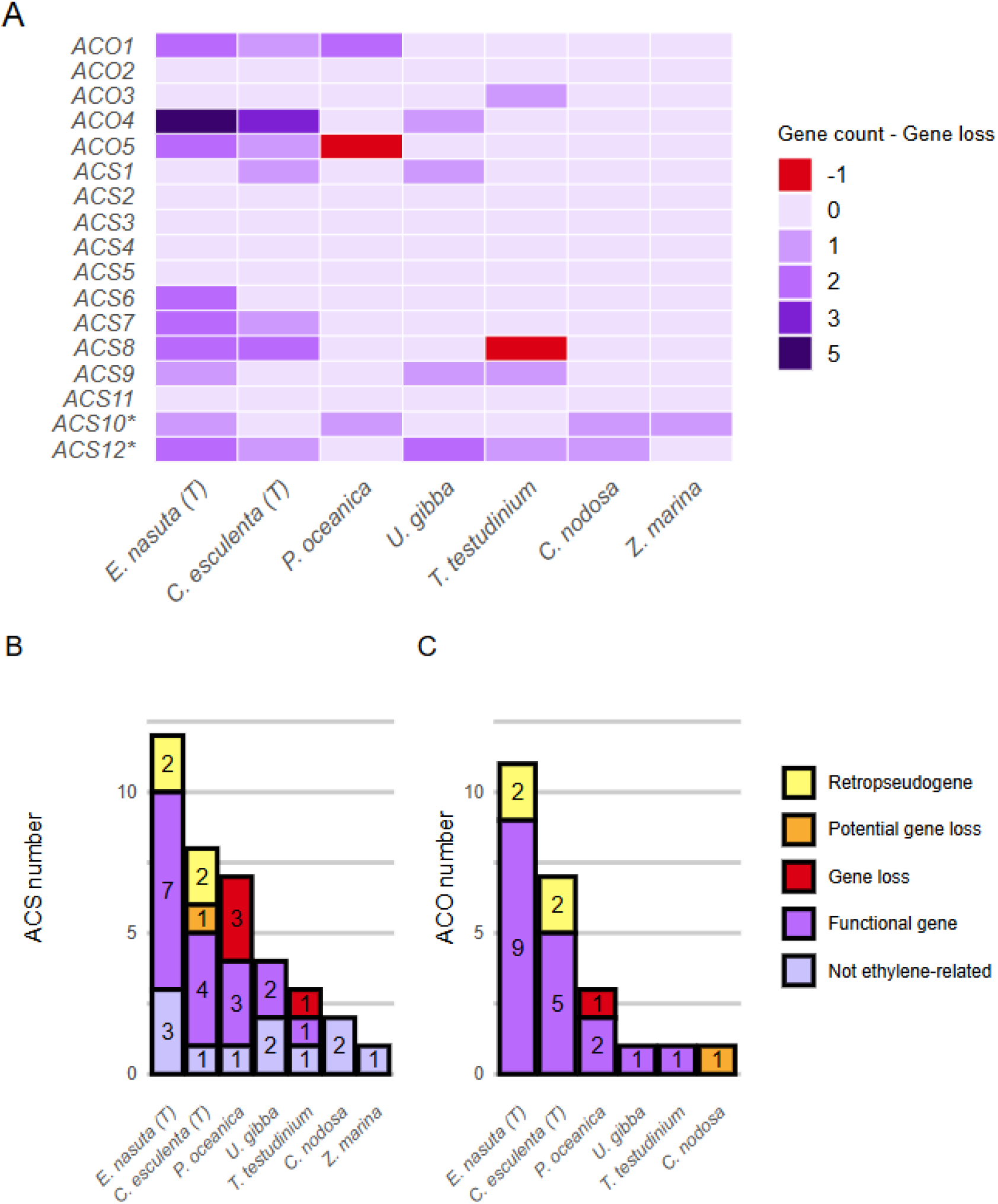
Comparison between the number of functional and non-functional *ACO* and *ACS* genes. A. Difference between the number of functional genes and the number of gene loss for all *ACO* and *ACS* genes, classified according to their homology with *A. thaliana* genes. * indicates genes that are probably not involved in ethylene synthesis. B. Overall results for *ACS* and C. Overall results for *ACO*. (T) indicates terrestrial species.

The only *ACO* or *ACS* genes found in *C. nodosa* and *Z. marina* are *ACS10* or *ACS12* genes, which are not involved in ethylene synthesis in *A. thaliana* (Yamagami et al., 2003). This confirms the loss of ethylene synthesis in these species. Surprisingly, we found no evidence of gene loss in these species; all the observed losses were found in *P. oceanica* and *T. testudnium*, the two marine species in which ethylene synthesis still exist. One explanation could be that *C. nodosa* and *Z. marina* lost the ability to generate ethylene at an early stage of their evolutionary history, to the point that even the pseudogenes have disappeared. In contrast, *P. oceanica* and *T. testudinium* may have a way of limiting ethylene accumulation, as suggested by Ma *et al* (Ma et al., 2024). Nevertheless, a decrease in the number of functional *ACO* and *ACS* genes is evident, and the finding of gene loss and unfragmented pseudogenes only in these species shows that the pseudogenization of these genes is a recent and ongoing process. By comparison, no strong evidence of *ACO* or *ACS* gene loss has been found in the land plant *E. nasuta* or the amphibian *C. esculenta*. Compared to *E. nasuta*, the submerged freshwater plant *U. gibba* appears to have undergone a strong reduction in the number of *ACS* and *ACO* genes, suggesting that this contraction is linked to an aquatic lifestyle and not solely to a marine one.

## Discussion

We were able to identify some *ACO* and *ACS* pseudogenes, likely originating from genes involved in ethylene synthesis, and confirmed the reduction in the number of these genes in seagrasses. In comparison, no evident gene loss was observed in terrestrial or amphibious plants. Our results thus confirm that the contraction of genes involved in ethylene biosynthesis in seagrass is a true contraction of these genes rather than an expansion in terrestrial plants, although a combination of both contraction and expansion remains possible. We also observed a reduction in the number of *ACO* and *ACS* genes in *U. gibba* compared to *E. nasuta*, suggesting that this reduction is linked to aquatic lifestyle, whether in fresh or salt water. However, we did not find any pseudogenes in *U. gibba, C. nodosa*, or *Z. marina*, even though the latter two have lost all of their *ACO* or *ACS* genes involved in ethylene synthesis. This suggests that the loss of these genes is more ancient in these species, whereas, conversely, all the predicted pseudogenes were found in *P. oceanica* and *T. testudinium*, showing that the loss of ethylene biosynthesis is an ongoing process. Our work highlights that pseudogenes can be used to confirm multigene family contractions and disappearing processes. However, we acknowledge that traces of loss of function seem to disappear rapidly, as no pseudogenes were found in species where ethylene biosynthesis has completely disappeared, and the total number of gene loss found remains small. Pseudogenes should therefore only be used for recent or ongoing loss of function.

## Methods

The genomes, protein sequences and annotations of the seagrasses were downloaded from https://bioinformatics.psb.ugent.be/gdb/seagrasses/ (Ma et al., 2024). Similar files for *C. esculenta, U. gibba* and *E. nastua* were respectively downloaded from CNGBdb (Yin et al., 2021), CoGe (Lan et al., 2017) and GenBank (ASM5117567v1). Scaffolds with a length < 300 kb were also removed, as well as any gene or protein related to those scaffolds. This was done to eliminate possible organelle genomes and small DNA fragments at the ends of which genes can be artificially “truncated” and annotated as pseudogenes. Finally, repeated sequences in the genomes were identified with Red v2.0 (Girgis, 2015) and hard-masked with bedtools v2.30 (Quinlan and Hall, 2010).

The functional annotation of the proteins was done using InterProScan v5.73 (Jones et al., 2014). To filter transposon proteins, a first set of transposon proteins was identified from all the protein sequences by their homology with transposable elements from the RepeatExplorer Viridiplantae v4.0 TE protein database (Novák et al., 2010). Homology was inferred using DIAMOND v2.1.9 (Buchfink et al., 2015) with --ultra-sensitive option. This set was then used to found other transposon proteins from all the protein sequences in a similar manner. Remaining sequences were further removed if they harbored any transposon-associated or virus-associated domain from CATH / Gene3D v4.4, (https://www.cathdb.info/) (Dawson et al., 2017), Pfam v38.1 (https://pfam.xfam.org/) (Bateman et al., 2004), PROSITE release 2026_01 (https://prosite.expasy.org/) (Sigrist et al., 2026), PANTHER v19.0 (https://pantherdb.org/) (Mi et al., 2005), SUPERFAMILY 2 (http://supfam.org) (Pandurangan et al., 2019) or Conserved Domain Database v3.21 (https://www.ncbi.nlm.nih.gov/Structure/cdd/cdd.shtml) (Wang et al., 2022) (Table S3). Pseudogenes structural annotation was done using P-GRe v1.0 (Cabanac et al., 2026) with options -A -Q and miniprot v0.18 dependency (Li, 2023). The transposon-free protein sequences from the different species were also concatenated and passed to the -u option to predict unitary pseudogenes. Functional annotations of the pseudogenes was then done by transferring annotations from their parent gene. R^2^ was calculated by squaring R’s cor() function output.

The functional *ACO* and *ACS* genes of the different species were found and categorized (*ACO1, ACO2, etc*.) according to their homology with the *ACO* and *ACS* genes of *A. thaliana* TAIR10 (Lamesch et al., 2012), downloaded from Ensembl Plants release 62 (https://plants.ensembl.org/index.html) (Bolser et al., 2016). Homology was inferred with DIAMOND v2.1.9 as described before. All the genes thus found were checked to identify whether they were in fact pseudogenes. The ACO and ACS proteins of *A. thaliana* were mapped around the genes that appeared truncated with miniprot v0.18, to retrieve any missing exon. The genes were thus re-classified as pseudogenes when no additional exons could be identified or when exons were identified but showed evidence of pseudogenization (frame-shifts or in-frame stop-codon). We also checked that every functional *ACS* genes had the conserved ACS-activity-related tripeptide Ser/Thr-Asn-Pro. All pseudogenes inferred either from the identified ACO or from ACS proteins were individually checked. Their reconstructed and frame-shift corrected protein sequences were used to determine which *A. thaliana* protein had the strongest similarity using DIAMOND v2.1.9. Pseudoproteins that did not correspond to ACO or ACS proteins were removed. It was also verified that the pseudogenes showed clear signs of pseudogenization (in-frame stop codon, frame shift, truncation, absence of a start or stop codon), and that they did not correspond to a missing part of an annotated functional gene by checking for the absence of *ACS* or *ACOs* genes within 20 kb of the pseudogene. Finally, we verified that the identified pseudogenes actually corresponded to gene loss events by checking that they had not been categorized as retrocopies (“unitary_pseudogene”) by P-GRe v1.0 (Cabanac, 2026). When the software failed to categorize them, we investigated deeper for possible intron losses by checking if the protein alignment performed by P-GRe v1.0 had split proteins between two amino acids that are normally encoded by a single exon.

## Acknowledgments

We are also thankful to the Toulouse University and CNRS for supporting our research.

